# Pseudomonas aeruginosa kills Staphylococcus aureus in a polyphosphate-dependent manner

**DOI:** 10.1101/2023.12.05.570291

**Authors:** Ritika Shah, Olivia Jankiewicz, Colton Johnson, Barry Livingston, Jan-Ulrik Dahl

## Abstract

Due to their frequent coexistence in many polymicrobial infections, including in patients with burn or chronic wounds or cystic fibrosis, recent studies have started to investigate the mechanistic details of the interaction between the opportunistic pathogens *Pseudomonas aeruginosa* and *Staphylococcus aureus*. *P. aeruginosa* rapidly outcompetes *S. aureus* under *in vitro* co-cultivation conditions, which is mediated by several of *P. aeruginosa*’s virulence factors. Here, we report that polyphosphate (polyP), an efficient stress defense system and virulence factor in *P. aeruginosa*, plays a role for the pathogen’s ability to inhibit and kill *S. aureus* in a contact-independent manner. We show that *P. aeruginosa* cells characterized by low polyP level are less detrimental to *S. aureus* growth and survival while the gram-positive pathogen is significantly more compromised by the presence of *P. aeruginosa* cells that produce high level of polyP. We show that the polyP-dependent phenotype could be a direct effect by the biopolymer, as polyP is present in the spent media and causes significant damage to the *S. aureus* cell envelope. However, more likely is that polyP’s effects are indirect through the regulation of one of *P. aeruginosa’s* virulence factors, pyocyanin. We show that pyocyanin production in *P. aeruginosa* occurs polyP-dependent and harms *S. aureus* through membrane damage and the generation of reactive oxygen species, resulting in increased expression of antioxidant enzymes. In summary, our study adds a new component to the list of biomolecules that the gram-negative pathogen *P. aeruginosa* generates to compete with *S. aureus* for resources.

**IMPORTANCE:** How do interactions between microorganisms shape the course of polymicrobial infections? Previous studies have provided evidence that the two opportunistic pathogens *P. aeruginosa* and *S. aureus* generate molecules that modulate their interaction with potentially significant impact on disease outcomes. Our study identified the biopolymer polyP as a new effector molecule that impacts *P. aeruginosa*’s interaction with *S. aureus*. We show that *P. aeruginosa* kills *S. aureus* in a polyP-dependent manner, which occurs primarily through the polyP-dependent production of the *P. aeruginosa* virulence factor pyocyanin. Our findings add a new role for polyP to an already extensive list of functions. A more in-depth understanding of how polyP influences interspecies interactions is critical, as targeting polyP synthesis in bacteria such as *P. aeruginosa* may have a significant impact on other microorganisms and potentially result in dynamic changes in the microbial composition.

## INTRODUCTION

Polymicrobial infections are characterized by the presence of microorganisms in the host that create a niche for additional microbial species to colonize and further impact pathogenesis (1). The interaction between two microorganisms can be beneficial to both or harmful to one and thus have significant consequences for the microbial composition at the infection site. While behavior and physiology of pathogens are often studied in single species, infections are often defined by their complex polymicrobial interactomes, which may have significant impacts on virulence, persistence, and/or antimicrobial tolerance of either one or several community members (2). Two of the most frequently co-isolated pathogens in polymicrobial infections are the gram-negative *P. aeruginosa* and the gram-positive *S. aureus,* which infect, among others, burn and chronic wounds as well as the lungs of cystic fibrosis (CF) patients (3). CF is an inheritable genetic disease in patients with defects in their CF transmembrane conductance regulator (CFTR) gene, affecting over 30,000 people only in the U.S. (4). The CF lungs get colonized by several bacterial species throughout life, and their composition changes as patients age (5). *S. aureus* is the most prevalent pathogen in lungs of CF patients and often already present at a very young age. However, over time, the *S. aureus* population declines and infections with *P. aeruginosa* become more prominent at a later stage in life, indicating a potential interaction between the two species (6). Likewise, wounds provide a favorable environment for polymicrobial infections involving both *S. aureus* and *P. aeruginosa,* which allow both pathogens to co-exist and mutually benefit from each other (7). Several independent studies provide clear evidence that the two species mutually affect their antimicrobial tolerance profiles making *S. aureus* / *P. aeruginosa* co-infections typically more severe and challenging to treat (8–10). However, while the underlying mechanisms are often poorly understood, they are likely attributed to the complex response systems that both species activate in co-culture (10). Both *S. aureus* and *P. aeruginosa* produce an array of virulence factors that often lead to invasive burn wound sepsis (11).

Due to its intrinsic and acquired resistance to almost all commonly used antibiotics, *P. aeruginosa* is one of the leading causes of hospital-acquired infections (12). This opportunistic pathogen produces a large army of virulence factors, such as 4-hydroxy-2-heptylquinoline-N-oxide (HQNO), elastase, rhamnolipids, pyocyanin, pyoverdine, and pyochelin, all of which contribute to the pathogenicity of this bacterium (13). Uncontrolled elastase production can lead to necrotic skin lesions and corneal ulcers in the host (14), while rhamnolipids are surfactant-like molecules that result in erythrocyte destruction and play a role for biofilm formation and protection (15). Pyochelin and pyoverdine serve as iron-chelating siderophores when *P. aeruginosa* experiences iron-limiting conditions (16). While both virulence factors were proposed to sequester iron from *S. aureus* cells that undergo cell lysis (17), pyochelin has also been shown to induce the production of hydroxyl radicals through the Fenton reaction, resulting in significant tissue damage (18). The fluorescent phenazine compound pyocyanin causes oxidative damage in host cells primarily by inducing the production of hydrogen peroxide while simultaneously reducing cellular glutathione level (19). Pyocyanin, which is present in sputum of CF patients and burn wounds at concentrations between 15-27 mg/ml (20), has also been shown to act as a potent inhibitor of aerobic respiration by targeting the electron transport chain (21). Much like *P. aeruginosa* does the gram-positive pathogen *S. aureus* have an intrinsic ability to resist multiple antibiotics and produce virulence factors that promote its ability to escape from innate immune cells (22). Several members of the *S. aureus* secretome have been identified as signals for *P. aeruginosa* when cultivated together (23, 24) and *S. aureus* has also evolved various strategies to evade killing by *P. aeruginosa,* such as the formation of small colony variants [recently reviewed in (17)].

Inorganic polyphosphate (polyP) is highly conserved, present in prokaryotic and eukaryotic species, and structurally extremely simple, as it consists of a linear arrangement of phosphate molecules (up to ∼1,000 Pi/chain) covalently linked via high-energy phosphoanhydride bonds (25, 26). In bacteria, generation of polyP_(n+1)_ is reversibly catalyzed by polyP kinase 1 (PPK1), an enzyme that transfers a terminal phosphate of ATP to a growing chain of polyP_(n)_ (27, 28). Degradation of polyP into monomeric Pi molecules is catalyzed by exopolyphosphatase (PPX) (28, 29). Gram-negative bacteria have long been known to produce long chains of polyP as a strategy to contribute to virulence, biofilm formation, motility, persister cell formation, and to counter host defense mechanisms (25, 30–33). PolyP has been identified as an important player for bacterial resistance towards host-generated antimicrobial oxidants: during periods of oxidative stress the cell converts most of its ATP pool into large amounts of polyP, which acts as chemical chaperone and binds to unfolded proteins to prevent aggregation (33, 34). When the stress subsides, polyP is degraded by PPX into inorganic phosphate molecules (25, 35). Due to the manifold functions of this ancient molecule, polyP-deficient strains display pleotropic phenotypes, including increased susceptibility to oxidative stress (33, 35), and defects in biofilm formation, motility, and several virulence factors such as elastase and rhamnolipids (25, 32, 36, 37). Moreover, deletion of PPK in the *P. aeruginosa* strain PAO1 resulted in increased susceptibility towards several different antibiotics, including ciprofloxacin, chloramphenicol, and rifampicin (38). Very similar results were obtained with other pathogens, including *Salmonella enterica, Vibrio cholera, and Mycobacterium tuberculosis*, where a PPK deletion significantly reduced bacterial pathogenicity as well as persister cell and biofilm formation (39–41). Independent studies confirmed that the presence of oxidative stress defense systems, including polyP, positively affect the pathogen colonization in the host, emphasizing the importance of polyP production for pathogenesis (42–44). Given that many medically relevant pathogens rely on polyP as protective mechanism, targeting bacteria-specific processes such as polyP production represent intriguing alternative treatment options, as polyP production only becomes essential for bacterial survival in the context of infections (12, 45). In fact, several PPK inhibitors have recently been discovered, which severely compromise bacterial survival during stress, biofilm-formation, colonization, and oxidant resistance (42, 44, 46).

In this study, we report that *P. aeruginosa*-generated polyP plays a role during competition with *S. aureus*. We show that *P. aeruginosa*-mediated inhibition and killing of *S. aureus* could occur directly through extracellular polyP. More likely, however, is an indirect effect, as polyP regulates the production of *P. aeruginosa* virulence factors, such as pyocyanin, which causes membrane damage and elicits oxidative stress in *S. aureus*.

## RESULTS

### *S. aureus* is inhibited by *P. aeruginosa* spent media in a polyP-dependent manner

Previous studies by others have made significant progress for our understanding of how the opportunistic pathogens *P. aeruginosa* and *S. aureus* affect each other, which likely has important implications on the trajectories of polymicrobial infections [recently reviewed in (17)]. Many gram-negative bacteria, such as *P. aeruginosa*, produce the biopolymer polyP when they experience various forms of stress, and polyP has also been shown to contribute to bacterial virulence (25, 30). To test whether *P. aeruginosa*’s ability to produce polyP has any impacts on the growth of *S. aureus*, we exposed the *S. aureus* strain USA300LAC to spent media of PA14 strains with defects in polyP metabolism that were cultivated to stationary phase for 24 hrs and sterile-filtered. We compared the inhibitory effects of spent media from cultures that are characterized by reduced cellular polyP level (*i.e. Δppk1* and *Δppk1:::*Gm^R^ strains, respectively) as well as dysfunctional polyP degradation (*i.e. Δppx* strain) with those from the corresponding wildtype PA14 by measuring the zones of inhibition of *S. aureus* growth in an agar cup assay after incubation for 16-18 hrs. The PA14Δ*lasI* strain was used as a positive control because disruption of *lasI* has previously been shown to result in reduced inhibition of *S. aureus* growth (47, 48). As a negative control, USA300LAC was exposed to spent media from its own 24 hrs culture, which had no inhibitory effect on the strain. As expected, we observed substantial zones of inhibition with a diameter of ∼28 mm in areas where the spent media of wild-type PA14 was spotted (**Fig. 1A**). The zones of inhibition were significantly reduced (∼17 mm diameter) when the spent media of *ΔlasI* cultures was spotted. Intriguingly, even though the spent media of the polyP-deficient mutants (*i.e. Δppk* and *Δppk:Gm^R^*) were able to inhibit *S. aureus*, the zones of inhibition were significantly smaller when compared to spent media from wild-type PA14 (i.e. ∼22 and ∼21 mm diameter, respectively, compared to ∼28 mm) (**Fig. 1A**). When USA300LAC was exposed to the *Δppx* spent media, the resulting zones of inhibition were much larger than those caused by PA14 wildtype (i.e. diameters of ∼33 mm compared to ∼28 mm) (**Figure 1A**).

**FIG 1.**
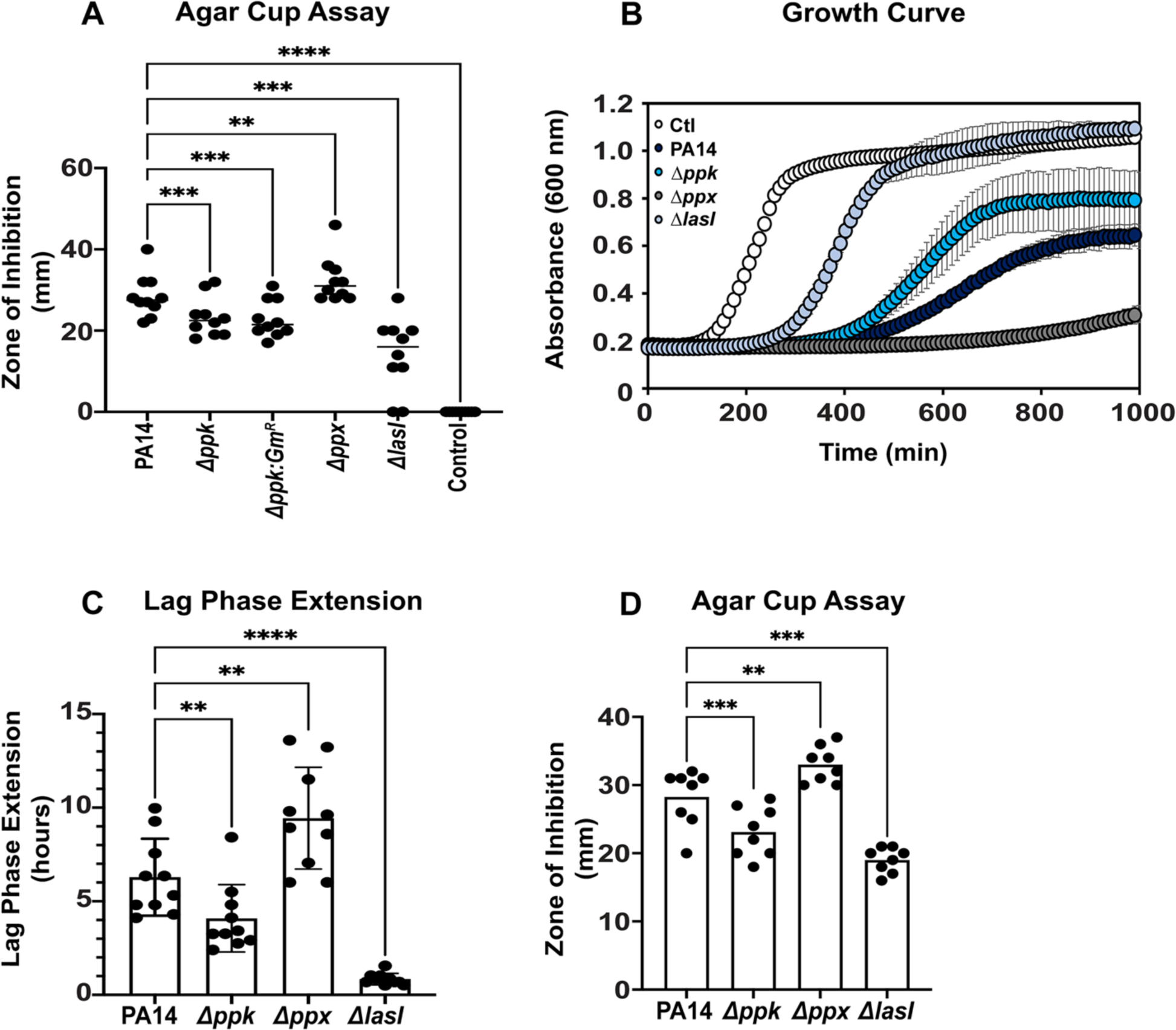
*S. aureus* is inhibited by *P. aeruginosa* spent media in a polyP-dependent manner. **(A)** *S. aureus* strain USA300LAC was diluted into LB media to an OD_600_ of 0.05 and spread onto LB agar. Wells were cut into the LB agar and filled with 150 µL spent media sterile filtered from the indicated 24 hrs, stationary phase *P. aeruginosa* cultures. The zones of inhibition were measured after incubation at 37 °C for 16-18 hrs *(n = 10, ± S.D.)*. **(B, C)** USA300LAC was diluted into TSB to an OD_600_ of 0.03 and exposed to the indicated *P. aeruginosa* spent media in a 15:1 (v/v) ratio. *A*_600nm_ was measured every 10 minutes for 16 hours using the Tecan Infinite 200 plate reader. White circles, control; dark blue circles, PA14; blue circles, Δ*ppk*; bright blue circles, Δ*lasI;* grey circles, Δ*ppx.* Lag phase extensions were calculated as described in *Material and Methods (n = 10, ± S.D.)* **(D)** *S. aureus* clinical isolate 93 was diluted into LB media to an OD_600_ of 0.05 and spread onto LB agar. Wells were cut into the LB agar and filled with 150 µL spent media of the indicated 24 hrs, stationary phase *P. aeruginosa* cultures. The zones of inhibition were measured after incubation at 37 °C for 16-18 hrs *(n = 8, ± S.D.). Statistical tests: One-Way ANOVA, ** p ≤ 0.01, *** p ≤ 0.001, **** p ≤ 0.0001*.

Next, we used a growth curve-based approach to verify the observed phenotypes from our agar cup assay. We exposed USA300LAC to sterile-filtered spent media of 24 hrs cultures of PA14, *Δppk, Δppx*, and *ΔlasI*, respectively, in a 15:1 (v/v) ratio and monitored growth over a time course of 16 hours. We and others previously reported that bacterial exposure to different stressors result in concentration-dependent extensions of the lag phase (LPE) (49–51). We quantified the LPEs for USA300LAC growth in the presence of the spent media of each of the different strains as described in the Materials and Methods section. Exposure of USA300LAC to spent media of wild-type PA14 caused substantial LPE (i.e. ∼6.3 hrs LPE) compared to USA300LAC cultures that were exposed to their own spent media (**Fig. 1B,C),** suggesting that PA14 spent media contains compounds inhibitory for *S. aureus* growth. Consistent with the results from our agar cup assay, the growth of USA300LAC was even more inhibited in the presence of spent media from the *Δppx* strain (i.e. ∼9.2 hrs LPE), whereas we observed the opposite effect when USA300LAC was treated with the sterile-filtered supernatant of the *Δppk* strain as evidenced by a less pronounced LPE compared to PA14 wildtype cultures (i.e. ∼3.8 hrs LPE) (**Fig. 1B,C**). As expected and reported before (52), spent media of the *ΔlasI* strain was least inhibitory also in the context of planktonic *S. aureus* growth (i.e. ∼1.2 hrs LPE) **(Fig. 1B,C)**. To exclude the possibility that the observed inhibition of *S. aureus* by *P. aeruginosa* is strain-dependent, we performed the agar cup assay using a *S. aureus* clinical isolate from a burn wound patient. We found that addition of the sterile-filtered supernatants of the PA14 strains wildtype, Δ*ppk*, Δ*ppx*, and Δ*lasI* resulted in a polyP-dependent inhibition of the *S. aureus* clinical isolate 193 **(Fig. 1D)** similar to what we had observed with USA300LAC **(Fig. 1A)**. In contrast, no growth inhibition was observed when we treated the different *P. aeruginosa* strains with spent media of stationary phase *S. aureus* USA300LAC cells **(Supplementary Fig. S1)**. Taken together, our results indicate that the spent media of stationary phase *P. aeruginosa* cells contains one or more inhibitory compounds that negatively affect *S. aureus* growth in a polyP-dependent manner, which could shape how the two pathogens interact with each other.

### *P. aeruginosa* kills *S. aureus* in a polyP-dependent manner

To examine whether *P. aeruginosa*’s polyP-dependent negative effect on *S. aureus* only affects growth or also survival, we performed a co-cultivation assay using actively growing USA300LAC cells in combination with any of the *P. aeruginosa* strains (*i.e.* PA14, *Δppk*, *Δppx*, and *ΔlasI,* respectively) at a 100:1 inoculation ratio. At the indicated time points, samples were serially diluted and plated onto mannitol salt agar to select for *S. aureus* and cetrimide agar for *P. aeruginosa* colonies, respectively. We confirmed in our co-cultivation experiment that PA14 wildtype indeed outcompetes *S. aureus* resulting in complete eradication of the Gram-positive pathogen after 21 hrs **(Fig. 2**, *dark blue circles*). The killing of *S. aureus* was even more pronounced in competition with *Δppx,* resulting in complete eradication after 18 hrs **(Fig. 2**, *grey circles*). On the other hand, *S. aureus* survival was significantly higher when USA300LAC was co-cultured with the polyP-deficient Δ*ppk* strain compared to the corresponding wildtype PA14, resulting in ∼ 2 log increased survival after 21 hrs **(Fig. 2**, *blue circles*). As expected, the competition between USA300LAC and the *lasI*-deficient PA14 strain had the least impact on *S. aureus* survival. Notably, the co-cultivation of the two pathogens had no effect on the survival of the different *P. aeruginosa* strains **(Supplementary Fig. S2)**. Overall, our data provide evidence that *P. aeruginosa* negatively impacts *S. aureus* survival in a contact-independent manner, which is mediated by *P. aeruginosa* polyP.

**FIG 2.**
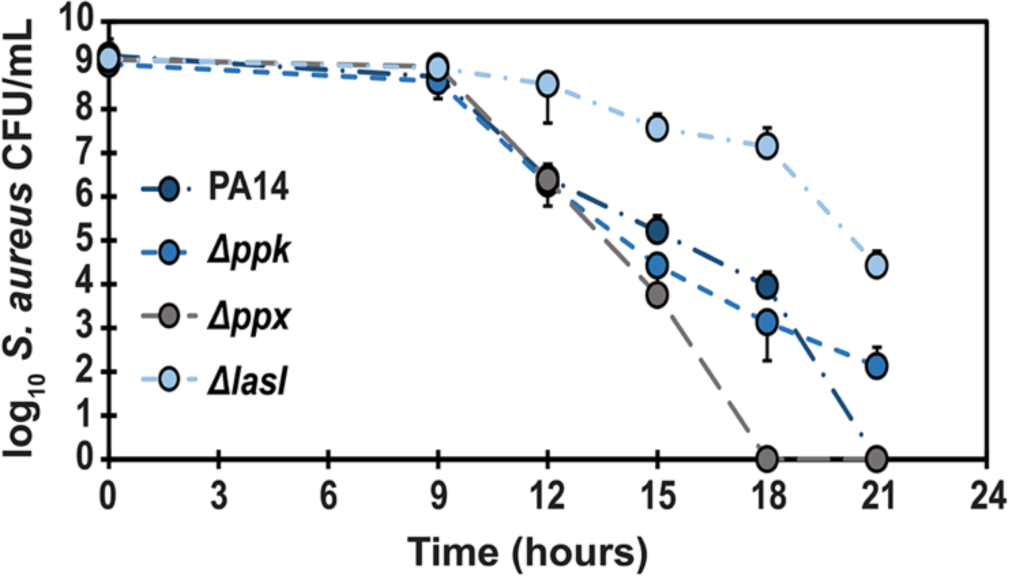
*P. aeruginosa* kills *S. aureus* in a polyP-dependent manner. Overnight cultures of *S. aureus* strain USA300LAC and the indicated *P. aeruginosa* strains were each diluted into TSB media to an OD_600_ of 1, mixed in a ratio of 100:1, and cultivated at 37 °C for 21 hrs. Samples were taken at indicated time points and serially diluted for *S. aureus* CFU counts on mannitol salt agar. Dark blue circles, PA14; blue circles, Δ*ppk*; bright blue circles, Δ*lasI;* grey circles, Δ*ppx; (n = 3, ± S.D.)*.

### PolyP contributes to *S. aureus* killing by causing membrane damage

Previous studies indicate that *S. aureus* is exceptionally sensitive to exogenous polyP and that gram-positive cell walls, such as those of *S. aureus*, can be significantly damaged by long-chain polyP, resulting in both bactericidal and bacteriolytic effects (53). Moreover, inactivation of the *P. aeruginosa ppk* gene has been linked to a substantial reduction in the production of some of *P. aeruginosa*’s most prominent virulence factors including biofilm formation, motility, elastase, and rhamnolipids (25, 30, 31). We therefore wondered whether the killing of *S. aureus* by *P. aeruginosa* occurs *(i)* directly through the release of polyP, *(ii)* indirectly through the secretion of one or more polyP-regulated *P. aeruginosa* virulence factors, or *(iii)* as a combination of the two. Given that the polyP-mediated effect of *P. aeruginosa* on *S. aureus* appears to appears to be contact independent (**FIG. 1**), we conclude that polyP must be present in the extracellular environment if it indeed contributes directly to the killing of *S. aureus*. Considering the average chain length of polyP in bacteria ranges from of 100-1000 orthophosphate molecules (54–56) and the fact that no polyP-specific export system has been identified yet make an active transport of polyP into the extracellular environment less likely. However, previous studies have shown that *P. aeruginosa* cells coordinate their group behavior through virulence factor-induced autolysis and the released virulence factors increase *P.* aeruginosa’s ability to form biofilms and increase resistance to beta-lactam antibiotics, which ultimately enables persistence of the population (57). Thus, differences in the extent to which cells lyse may be affected by polyP. We utilized an established assay that detects polyP by DAPI fluorescence (58) to quantify extracellular polyP level in the spent media of 24 hrs, stationary phase cultures of the PA14 wildtype, Δ*ppk*, Δ*ppx,* and Δ*lasI* strains. To our surprise, the spent media of all four PA14 strains contained very similar polyP level **(FIG 3A)**. *P. aeruginosa* autolysis also results in the release of DNA. We then measured DNA level in each of the different spent media and saw a very similar trend **(Supplementary FIG S3A)**, suggesting a comparable degree of cell lysis between all four strains. Despite our observation that overnight cultures of these strains differ in their optical density at 600 nm (data not shown), colony forming units (CFU) counts after 24 and 38 hrs, respectively, did not result in any significant differences **(Supplementary FIG S3B&C)**. To examine the degree to which the spent media of PA14 strains with defects in polyP metabolism cause membrane damage in *S. aureus*, we first confirmed that addition of exogenous polyP to actively growing *S. aureus* cells indeed results in increased propidium iodide (PI) fluorescence, a common method for detection of membrane damage **(FIG 3B)**. Non-specific binding of PI to nucleic acids enhances PI fluorescence exponentially, however, due to its size and charge, PI cannot cross intact inner membranes and therefore PI fluorescence correlates with the degree of membrane damage (59). We exposed USA300LAC to the 24 hrs spent media of the different PA14 strains and quantified their PI fluorescence after 1 hr of incubation. Intriguingly, PI fluorescence values of *S. aureus* cells exposed to the spent media of the *Δppk* and *ΔlasI* strains were significantly lower compared to PA14 wildtype spent media, whereas we did not observe any significant differences between PA14 wildtype and the polyP-accumulating strain *Δppx* **(FIG 3C)**. The membrane damaging effects of polyP directly affect *S. aureus* growth and survival: when actively growing USA300LAC cells were exposed to increasing concentrations of exogenous sodium polyP, we observed a concentration-dependent increase in the generation time as well as decrease in growth yield (**FIG 3D**) and a decline in CFU after 24 hrs of incubation (**Figure 3E**). Taken together, these results show that while polyP has a detrimental effect on the growth and survival of *S. aureus*, it likely does not play a major direct role in the *P. aeruginosa* polyP-mediated inhibition and killing of *S. aureus*.

**FIG 3.**
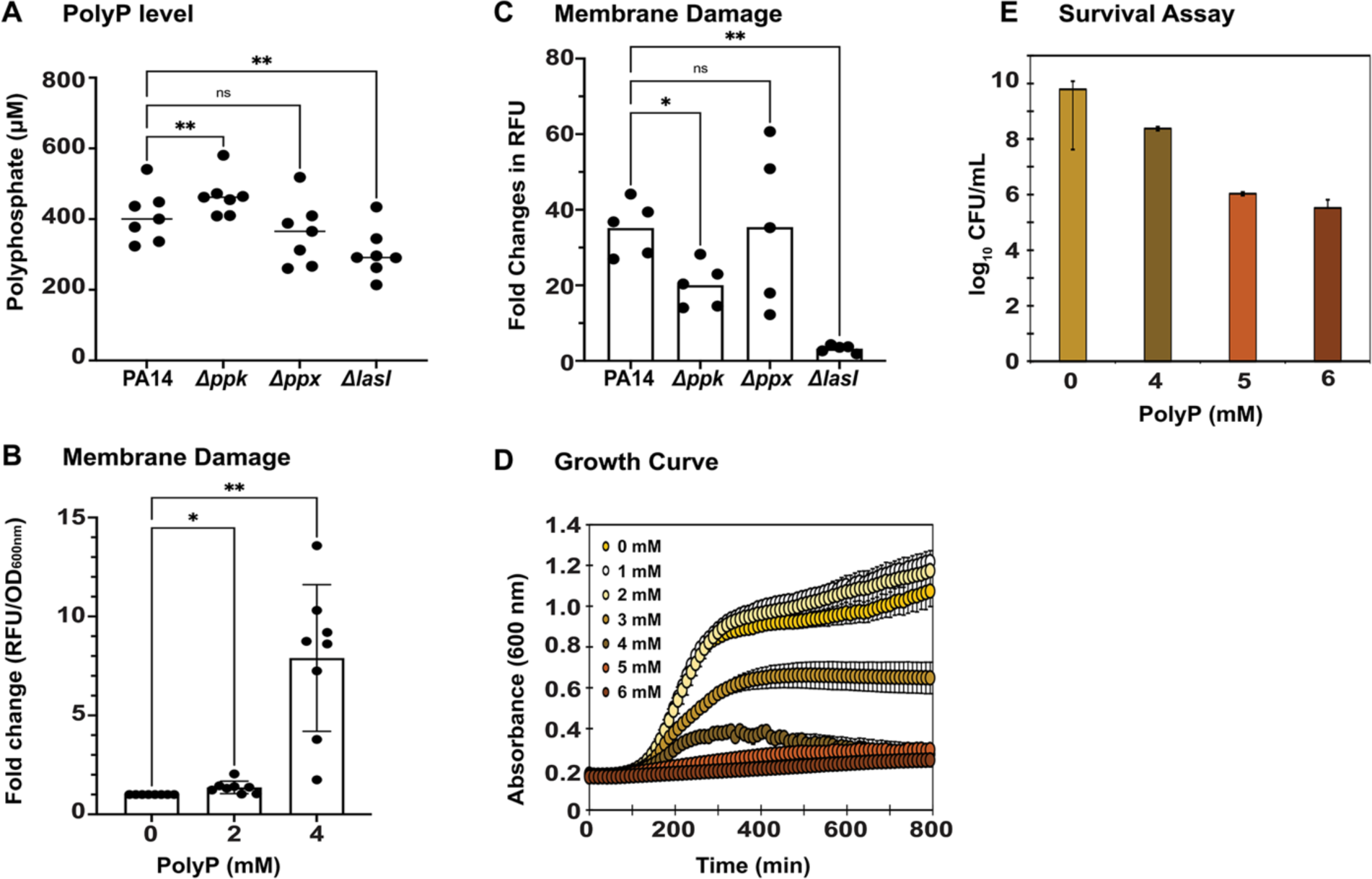
**PolyP contributes to *S. aureus* killing by causing membrane damage. (A**) PolyP level were determined in the spent media of 24 hrs, stationary phase *P. aeruginosa* cultures using 25 µM DAPI. DAPI-polyP fluorescence was measured at excitation/emission wavelengths of 415 and 550 nm, respectively. PolyP concentrations were calculated using a sodium polyP standard curve; *(n = 7, ± S.D.).* **(B)** Exponentially growing USA300LAC (OD_600_: 0.2) was exposed to the indicated concentrations of sodium polyP for 1 hour at 37 ^0^C, following which membrane damage was analyzed using 0.5 µM propidium iodide *(n = 8, ± S.D.)*. **(C)** Exponentially growing USA300LAC (OD_600_: 0.5) was exposed to spent media of the different *P. aeruginosa* strains for 1 hour at 37 ^0^C, following which membrane damage was analyzed using 0.5 µM propidium iodide *(n = 5, ± S.D.)*. **(D)** USA300LAC was diluted into TSB to an OD_600_ of 0.03 and exposed to increasing concentrations of sodium polyP. *A*_600nm_ was measured every 10 minutes for 16 hours using the Tecan Infinite 200 plate reader. *(n = 5 [with 4 technical replicates], ± S.D.)*. **(E)** USA300LAC was diluted into TSB to an OD_600_ of 0.03 and exposed to increasing concentrations of sodium polyP. USA300LAC survival was determined through CFU counts on TSB agar plates; *(n = 3, ± S.D.)*. *Statistical tests: One-Way ANOVA, ns: p > 0.05, * p ≤ 0.05, ** p ≤ 0.01*.

### *S. aureus* killing is alleviated in the absence of *P. aeruginosa* virulence factors

*P. aeruginosa* produces a large arsenal of virulence factors such as elastase, proteases, pyochelin, pyocyanin, pyoverdine, and rhamnolipids, all of which are secreted into the extracellular environment (60–62). Next, we tested whether and to which extent these virulence factors, at endogenous concentrations present in stationary phase PA14 cultures, affect the growth of *S. aureus* strain USA300LAC. We therefore compared the growth of USA300LAC in the presence and absence of spent media of strains with defects pyocyanin (i.e. *ΔphzM*), pyoverdine (i.e. *ΔpvdS*), rhamnolipids (i.e. *ΔrhlA*), elastase (i.e. *ΔlasB*), and pyochelin (i.e. *ΔpchB*) production to treatment with spent media of PA14 wildtype cells. We found that exposure to spent media of any of the five transposon mutant strains was significantly less inhibitory to USA300LAC growth compared to spent media of the corresponding wildtype PA14 **(FIG 4)** indicating that each of the virulence factors tested may contribute to *P. aeruginosa*-mediated inhibition of *S. aureus*. Our next goal was to determine whether the production of one or more of these virulence factors is regulated by polyP. We therefore extracted and quantified each virulence factor from the spent media of the *P. aeruginosa* PA14 strains wildtype, *Δppk*, *Δppx*, *ΔlasI*, and the transposon insertion mutant that lacks the ability to produce the respective virulence factor. The level of elastase **(Supplementary FIG 4A)**, LasA protease **(Supplementary FIG 4B)**, pyochelin **(Supplementary FIG 4C)**, pyoverdine **(Supplementary FIG 4D)**, and rhamnolipids **(Supplementary FIG 4E)** from the spent media of strains with defects in polyP metabolism revealed no significant differences suggesting that their production may not be controlled by polyP under the conditions tested.

**FIG 4.**
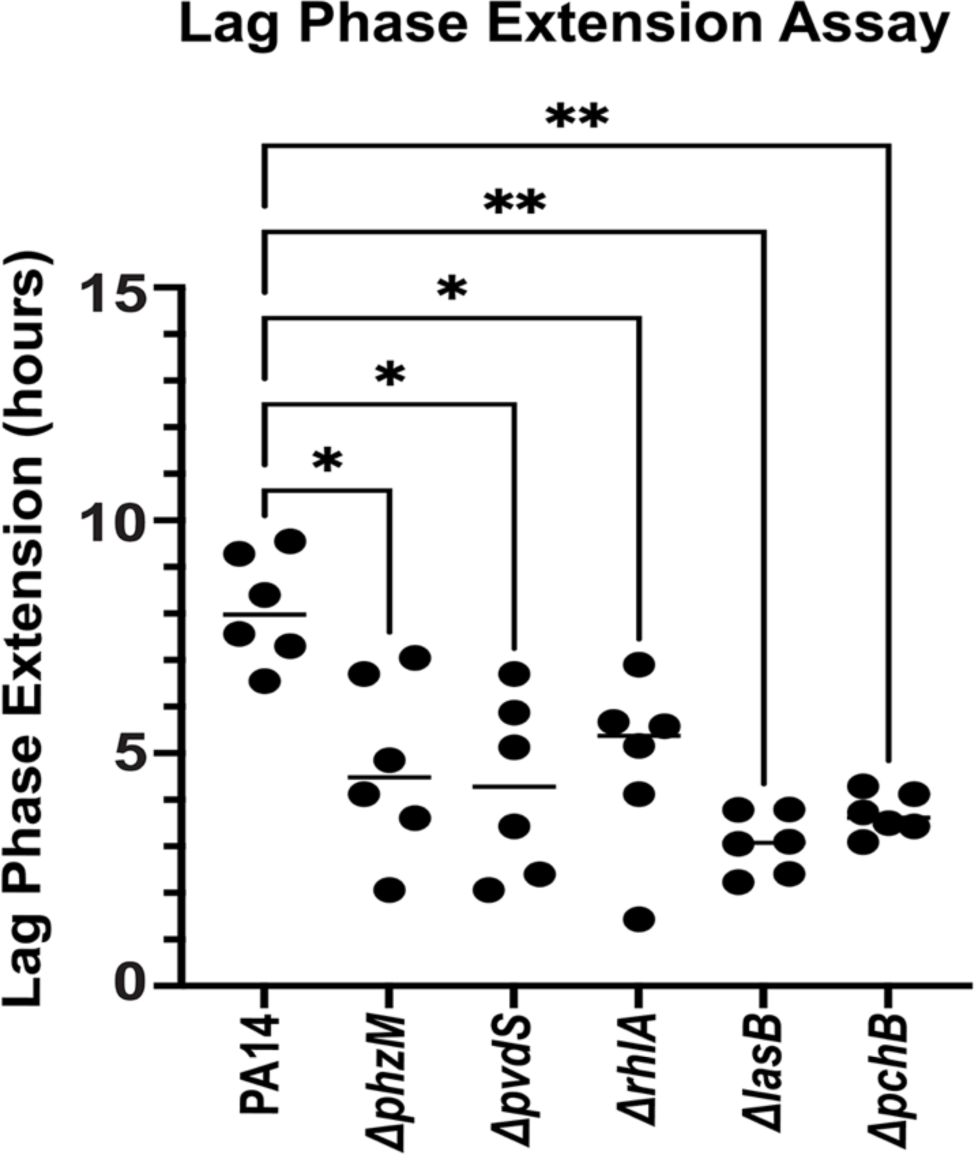
*S. aureus* killing is alleviated in the absence of *P. aeruginosa* virulence factors. *S. aureus* strain USA300LAC was diluted into TSB to an OD_600_ of 0.03 and exposed to the indicated *P. aeruginosa* spent media in a 15:1 (v/v) ratio. *A*_600nm_ was measured every 10 minutes for 16 hours using the Tecan Infinite 200 plate reader. Lag phase extensions were calculated as described in *Material and Methods; n = 6, ± S.D.; One-Way ANOVA, * p ≤ 0.05, ** p ≤ 0.01*.

### PolyP-mediated differences in pyocyanin level may contribute to the killing of *S. aureus* through the induction of oxidative stress and membrane damage

Exposure to spent media of the *ΔphzM* strain, which completely lacks pyocyanin production (63, 64), was also less inhibitory to *S. aureus* in our growth curve-based assay **(FIG 4)** suggesting that its absence reduces the susceptibility of *S. aureus* to *P. aeruginosa*. However, in contrast to the other virulence factors tested, quantification of pyocyanin level from the spent media of the different *P. aeruginosa* strains showed significant differences for the strains with defects in polyP metabolism: compared to the PA14 wildtype, pyocyanin level were significantly higher in cultures with increased cellular polyP level (i.e., *Δppx*) and substantially lower in cells with compromised polyP production (i.e., *Δppk*) (**FIG 5A**). Thus, our data support the possibility that polyP regulation of pyocyanin could play a role for the polyP-dependent inhibition of *S. aureus* by *P. aeruginosa.* To test whether *S. aureus* is indeed susceptible to pyocyanin, we exposed *S. aureus* to increasing concentrations of exogenous pyocyanin and monitored the growth in our lag phase extension (LPE) assay. Intriguingly, starting with as little as 5 mg/ml pyocyanin, increasing concentrations of pyocyanin elicit concentration dependent LPE in *S. aureus* **(FIG 5B)**. Exposure of *S. aureus* to increasing concentrations of pyocyanin resulted in a concentration-dependent decrease in survival **(FIG 5C)**. To investigate whether pyocyanin inhibits *S. aureus* growth and survival by damaging its cell membrane, we exposed exponentially growing USA300LAC cells to increasing concentrations of exogenous pyocyanin and determined PI fluorescence as a proxy for membrane damage. *S. aureus* cells that were incubated with 5 and 10 mg/ml pyocyanin for 1 hr showed a 1.8- and 2.8-fold increase in PI fluorescence compared to the untreated controls, suggesting that already low concentrations of pyocyanin have profound effects on the membrane integrity of *S. aureus* **(FIG 5D)**. Pyocyanin has long been shown to have damaging effects on human cells through the production of ROS, in particular hydrogen peroxide (65). We then investigated whether exposure to the spent media of the different *P. aeruginosa* strains would elicit an oxidative stress response in *S. aureus* in a polyP and pyocyanin dependent manner. We exposed *S. aureus* for 10 minutes to the different *P. aeruginosa* spent media in a 3:1 (v/v) ratio and determined changes in *katB* transcript level (*katB* encodes catalase B) using quantitative real-time PCR (qRT-PCR). Given the increased oxidative stress resistance of most *S. aureus* wildtype strains, we performed this experiment in USA300JE2*ΔkatAΔahpC*, a catalase A (*katA*) and peroxidase (*ahpC*) deficient strain that is characterized by significantly higher susceptibility to hydrogen peroxide **(Supplementary FIG S5)**. Compared to control samples, where exponentially growing USA300LAC was exposed to spent media from its own 24 hrs culture, treatment with spent media of the PA14 wildtype caused a ∼8-fold increase in *katB* transcript level **(FIG 5E)**. *katB* transcript level were significantly lower in the presence of the spent media of the *Δppk* and slightly but not statistically significantly increased upon exposure to the *Δppx* spent media **(FIG 5E)**. We observed almost no induction in *katB* mRNA level when *S. aureus* was treated with the spent media of the pyocyanin-deficient Δ*phzM* strain, whereas exogenous addition of the virulence factor induced *katB* transcription approximately 6-fold **(FIG 5E)**. Taken together, our data reveal that *P. aeruginosa*’s ability to produce pyocyanin is regulated by polyP. Pyocyanin negatively affects *S. aureus* growth and survival by causing membrane damage and inducing oxidative stress.

**FIG 5.**
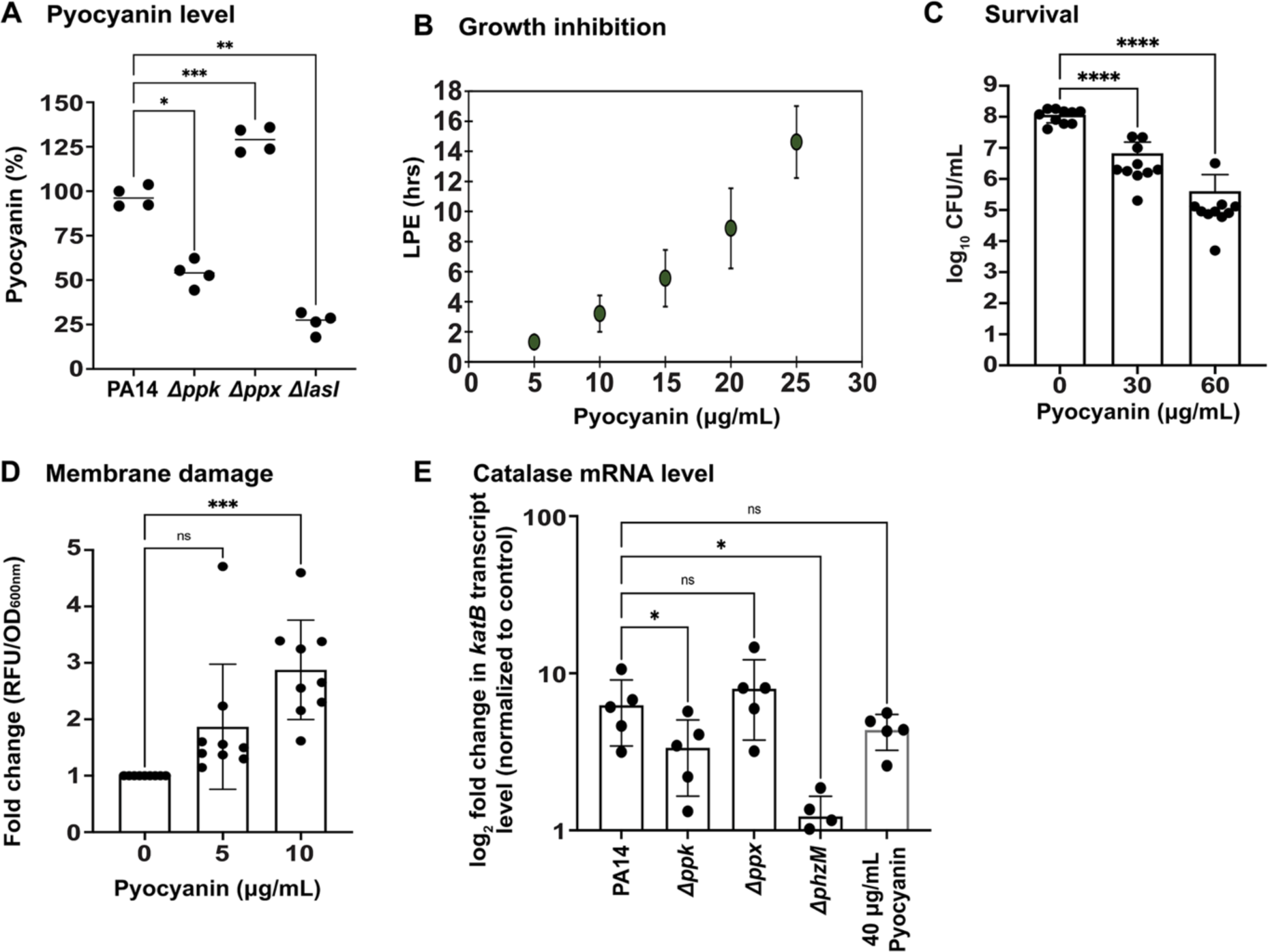
PolyP-mediated differences in pyocyanin level may contribute to the killing of *S. aureus* through the induction of oxidative stress and membrane damage. **(A)** Pyocyanin was quantified from the sterile-filtered spent media of the different 24 hrs *P. aeruginosa* cultures, *(n = 4, ± S.D.)*. **(B)** *S. aureus* strain USA300LAC was diluted into TSB to an OD_600_ of 0.03 and cultivated in the presence and absence of the indicated pyocyanin concentrations. *A*_600nm_ was measured every 10 minutes for 16 hours using the Tecan Infinite 200 plate reader. Lag phase extensions were calculated as described in *Material and Methods; (n = 3, ± S.D.).* **(C)** *S. aureus* strain USA300LAC was diluted into TSB to an OD_600_ of 0.05 and exposed to increasing concentrations of pyocyanin. USA300LAC survival was determined through CFU counts on TSA plates*; (n = 10, ± S.D.).* **(D)** Exponentially growing USA300LAC (OD_600_: 0.2) was exposed to the indicated concentrations of pyocyanin for 1 hour at 37 ^0^C, following which membrane damage was analyzed using 0.5 µM propidium iodide; *(n = 9, ± S.D.)* **(E)** Exponentially growing USA300JE2*ΔkatAΔahp* were exposed for 10 min to sterile-filtered spent media of the indicated PA14 cultures in a 3:1 (v/v) ratio or to 40 µg/ml pyocyanin, respectively. Induction of *katB* transcript level was determined using qRT-PCR; *(n = 4, ± S.D.).* Statistical tests: *One-Way ANOVA, ns: p > 0.05, * p ≤ 0.05, ** p ≤ 0.01, *** p ≤ 0.001)*.

## DISCUSSION

In the present study, we investigated the role of the bacterial stress defense system polyP for the interaction between the opportunistic pathogens *P. aeruginosa* and *S. aureus* during *in vitro* co-cultivation. Previous work by others provided strong evidence that *P. aeruginosa* quickly outcompetes *S. aureus* under both planktonic and biofilm growth conditions (17, 66–69). The goal of our study was to identify additional factors that *P. aeruginosa* employs to inhibit and kill *S. aureus*. Our data revealed that the extent to which *S. aureus* is eradicated correlates with *P. aeruginosa’s* ability to produce polyP. Based on our findings, we propose the following model, which considers both direct and indirect contributions by polyP **(FIG 6)**. When *P. aeruginosa* cells undergo cell lysis polyP is released into the extra cellular environment and harms *S. aureus* directly by inducing membrane damage. Indirect effects of polyP on *S. aureus* are mediated by pyocyanin, a *P. aeruginosa* virulence factor produced in a polyP-dependent manner. *P. aeruginosa* secretes pyocyanin, which harms *S. aureus* in at least two distinct ways: *(i)* by targeting the inner membrane of *S. aureus* cells negatively affecting the membrane integrity of the gram-positive pathogen; and *(ii)* by inducing oxidative stress through the production of ROS, such hydrogen peroxide and superoxide radicals, which *S. aureus* compensates by upregulating the expression of antioxidant enzymes.

**FIG 6.**
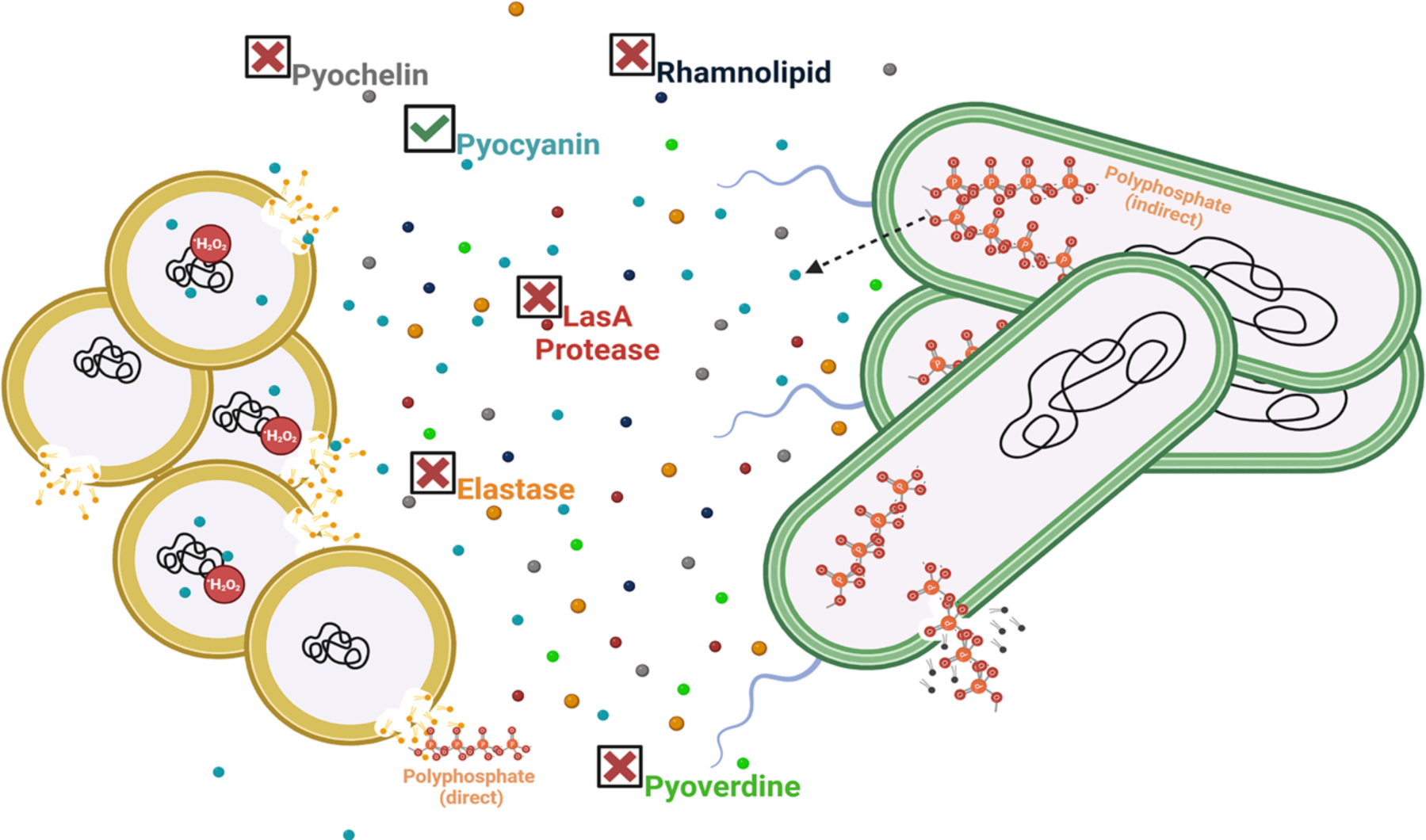
Current model of polyP’s role during *P. aeruginosa*-mediated killing of *S. aureus*. *P. aeruginosa* (green cells) sense *S. aureus* (yellow cells) and generate polyP and several virulence factors including pyochelin, rhamnolipid, pyocyanin, pyoverdine, elastase, and LasA, which are released into the extracellular environment. The release of polyP likely occurs through autolysis of *P. aeruginosa* and can cause damage to the *S. aureus* cell envelope, negatively impacting survival. However, polyP regulates the production of the *P. aeruginosa* virulence factor pyocyanin, which causes significant membrane damage and elicits the increased levels of antioxidant enzymes indicating oxidative stress.

The structurally very simple biopolymer polyP is composed of long chains of phosphate molecules linked via phosphoanhydride bonds and represents one of the most versatile posttranslational response systems that many gram-negative and some gram-positive bacteria generate during environmental stress (25). While polyP has been detected in both *P. aeruginosa* and *S. aureus* (44, 70) the enzymes required for polyP metabolism have only been identified in *P. aeruginosa*: the non-essential polyP kinase 1 (PPK1) transfers the terminal phosphate of ATP onto a growing chain of polyP_(n)_ polyP_(n+1)_ (27, 28, 71) whereas polyP is degraded into monomeric phosphate molecules by the exopolyphosphatase PPX (28). Thus, *ppk1*-deficient strains are typically characterized by very low amounts of endogenous polyP compared to the corresponding wildtype cells whereas strains lacking PPX display increased levels of polyP for a longer period of time (35). Several stressors have been identified that stimulate the production and cellular accumulation of polyP, including growth in stationary phase and during nutrient starvation (72, 73). Moreover, exposure to reactive oxygen and chlorine species (34, 74), osmotic imbalances, low pH (75), and elevated temperatures (76) induce polyP production. It is therefore not surprising that *ppk1*-deficient cells are characterized by pleiotropic phenotypes, such as increased susceptibility towards oxidative stress (33, 34, 77), antibiotics (78), elevated temperatures (76, 77), heavy metals (25), lipid A modification (78), and starvation (78). In addition, various studies point towards high sensitivity of bacteria towards changes in polyP metabolism given that both a reduction in the cellular polyP level (i.e. in the absence of the polyP-generating enzyme PPK or in the presence of PPK inhibitors) as well as polyP accumulation (i.e. in the absence of the polyP-degradation enzyme PPX) result in changes in motility and biofilm formation as well as attenuated production of virulence factors and colibactin, which are associated with effects on acute and chronic infections (30, 36, 79) Our present study adds a new phenotype for cells with defects in polyP metabolism to a growing list; the extent to which *P. aeruginosa* kills its competitor *S. aureus* correlates with the ability of the gram-negative opportunistic pathogen to produce and accumulate polyP (**FIG 2**). We conclude that the polyP-dependent inhibition of *S. aureus* by *P. aeruginosa* does not require a direct contact between the two pathogens as the sterile spent media of *P. aeruginosa* strains with defects in polyP metabolism was sufficient to cause a phenotypically similar growth inhibition of *S. aureus* (**FIG 1**). Our data suggest that *S. aureus* killing by *P. aeruginosa* is mediated by one or more inhibitory exoproducts that are either secreted into the extracellular environment or released by cell lysis or an otherwise unknown mechanism. It is well established that *P. aeruginosa* cells coordinate their group behavior through virulence factor-induced autolysis (57). Specifically, HQNO disrupts the electron transport chain, which yields in increased reduction of molecular oxygen and intracellular ROS accumulation. As a result, cells experience significant membrane damage, undergo autolysis, and release many cytoplasmic components, such as polyP and DNA, into the extracellular environment, where they facilitate biofilm formation and increase resistance to beta-lactam antibiotics (57). While there has not yet been identified a mechanism for the active secretion of polyP in *P. aeruginosa*, a recent study by Rijal *et al.* reported that *Mycobacterium* species secrete polyP during phagocytosis in macrophages and *Dictyostelium discoideum*, which significantly increases their chances for survival (80). Likewise, exogenous addition of long-chain polyP, as it is produced in bacteria, has been shown to increase bacterial survival in the phagosome. In our study, we found that the addition of exogenous polyP to actively growing *S. aureus* cells is detrimental to their growth and survival in a concentration-dependent manner (**FIG 3 D,E**). This effect is likely a consequence of the membrane damage that polyP elicits (**FIG 3B**). Due to its antimicrobial properties, polyP has long been used in the food industry to prevent food spoilage by gram-positive organisms ((53). When present at sufficiently high concentrations, polyP disrupts the bacterial cell envelope by chelating divalent cations from the membrane ((53, 81). Similarly, the spent media of *P. aeruginosa* strains characterized by either impaired polyP production or defective polyP degradation caused membrane damage to an extent that correlated with the intracellular polyP level (**FIG 3C**). However, the amounts of extracellular polyP and DNA in the spent media were not significantly different in these strains (**FIG 3A**) suggesting that the differences in *S. aureus* killing can likely not be attributed to polyP directly.

*P. aeruginosa* produces a plethora of virulence factors that are detrimental for *S. aureus* growth and survival, including respiratory toxins (i.e. hydrogen cyanide and HQNO), proteases (i.e. LasA, LasB), siderophores (i.e. pyoverdine, pyochelin), rhamnolipids, and N-acyl homoserine lactones (17), providing an alternative explanation for the polyP-dependent killing of *S. aureus*. PolyP has been shown to regulate the production of some of these virulence factors, including elastase LasB and rhamnolipids, although this had only been examined on transcriptional level (36). Quantification of elastase and rhamnolipids in the spent media of *P. aeruginosa* strains with different defects in polyP metabolism did not reveal any significant differences, suggesting that their production is not controlled by polyP under our experimental conditions, and therefore may not explain the phenotypic differences observed in our growth and survival studies. Likewise, pyochelin level and LasA protease activity showed no significant differences in polyP-compromised mutants compared to the wildtype suggesting that polyP is not involved in regulating their production. Intriguingly, we saw a significant drop in pyoverdine level in *P. aeruginosa* cells with defects in polyP production (i.e. Δ*ppk*). Pyoverdine is a major siderophore and involved in iron acquisition, which supports the formation of *P. aeruginosa* biofilms and contributes to virulence (82, 83). Previous studies have shown that *P. aeruginosa* secretes pyoverdine to bind iron from other competing microorganisms, including *S. aureus* strains that undergo cell lysis (19, 67). However, *P. aeruginosa* generates large amounts of the redox active phenazine molecule pyocyanin, production of which appears to be controlled by polyP: *P. aeruginosa* cells with defects in polyP production (i.e. Δ*ppk*) were characterized by a ∼50% reduction in pyocyanin compared to the corresponding wildtype, whereas the levels were significantly increased in Δ*ppx* cells, which accumulate polyP (**FIG 5A**). Pyocyanin is known to affect other commensal microbiota and be harmful to host cells, as the virulence factor inhibits aerobic respiration and elicits ROS production (17, 84). Our study revealed that *S. aureus* is extremely sensitive to exogenously added pyocyanin at physiologically relevant concentrations (20) resulting in significant growth inhibition **(FIG 5B)** and killing **(FIG 5C)** likely due to membrane damage (**FIG 5D**). Our results are in line with previous studies that were conducted to better understand pyocyanin’s role when *P. aeruginosa* competes with other pathogens and establishes infections (19, 85–87). We therefore attribute the differences in membrane damage, which we observed when *S. aureus* was exposed to the different spent media, to the varying extracellular concentrations of pyocyanin (**FIG 3C**). Consistent with these findings are our data that show efficient inhibition of *S. aureus* relies on functional pyocyanin production given that the spent media of the pyocyanin-deficient strain (i.e. Δ*phzM*) was significantly less inhibitory than the corresponding wildtype (**FIG 4**). Whether the membrane-damaging effects of pyocyanin is direct or indirect is still unclear but it has been proposed to be a consequence of pyocyanin-induced ROS production. Indeed, exposure to exogenously added pyocyanin caused a significant upregulation of *katB* transcript levels **(FIG 5E)**, one of the major catalases present in *S. aureus* that detoxifies hydrogen peroxide, suggesting that pyocyanin also induces ROS levels in *S. aureus*. Intriguingly, we observed differences in *katB* transcript levels of *S. aureus* exposed to the different spent media that correlated with their polyP and pyocyanin level **(FIG 5E)**. Thus, we provide novel insights into the role pyocyanin plays for *P. aeruginosa* mediated killing of *S. aureus*, which is regulated by polyP and causes oxidative stress in *S. aureus*.

Given the manifold phenotypes of Δ*ppk1* strains that range from defects in bacterial virulence, motility, quorum sensing, biofilm formation, to increased susceptibility to many environmental stressors [recently reviewed in (32, 33, 71, 88, 89)], it is not surprising that the polyP synthesizing enzyme PPK was revisited as a novel antimicrobial drug target. Several inhibitors of polyP production have been identified and characterized, such as the anti-inflammatory drug mesalamine (46), ellagic acid isolated from *Terminalia chebula* (90), and gallein (44). The inhibitory effects of mesalamine and gallein on polyP production was not limited to *P. aeruginosa* but also shown in many other bacterial pathogens, including uropathogenic *Escherichia coli*, *Vibrio cholerae*, *Klebsiella pneumoniae* and *Acinetobacter baumannii* (34, 91). Our work provides novel insights into the role of polyP for interspecies interactions and advises caution with regards to the use of PPK inhibitors in polymicrobial infection as this could have significant effects on other community members and e.g. result in an increased survival of *S. aureus,* a difficult to treat pathogen capable of infecting various organs and tissues.

## MATERIAL & METHODS

### Strains and growth conditions

All strains used in this study are listed in Table S1. The PA14 non-redundant transposon insertion mutant strains were used from the commercially available library described in (92). The clean deletion of the *Δppk* strain was constructed as described previously (34). All *P. aeruginosa* strains were cultivated in Luria Bertani (LB) broth at 37 ^0^C and 300 rpm shaking conditions. In contrast, overnight cultures of *S. aureus* were grown in LB broth and then transferred into Tryptic Soy Broth (TSB) to study inhibition by *P. aeruginosa* cells and sterile spent media, respectively. Gentamycin (15 µg/mL) was added whenever required.

### Collection of spent media

5 mL LB broth was inoculated with 50 ml of the respective overnight culture, incubated at 37 ^0^C and 300 rpm for 24 hours, and centrifuged at 4,255 x g for 15 minutes at 4 ^0^C. The supernatant/spent media was sterile filtered using a 0.22 µM filter and either used freshly or stored at -20 ^0^C.

### Determining inhibition of *S. aureus* by *P. aeruginosa* spent media using agar cup assays

The experiment was performed as previously described with slight modifications (93). Briefly, overnight *S. aureus* cultures were diluted to an OD_600_ of 0.05 and spread as a lawn onto LB agar. 0.05 cm wells were punctured into the agar and filled with 150 µL of sterile-filtered spent media from the indicated *P. aeruginosa* strains. The plates were then incubated overnight at 37 ^0^C to measure the respective zones of inhibition.

### Determining inhibition of *S. aureus* by *P. aeruginosa* spent media using a growth curve-based assay

*S. aureus* strains were grown overnight in LB, diluted into TSB to an OD_600_ of 0.02, and added to sterile-filtered spent media of different *P. aeruginosa* strains in the indicated ratios. Cells were then incubated with shaking at 37 ^0^C and OD_600_ measurements taken every 10 minutes for 16 hours in a Tecan Infinite 200 plate reader. The sensitivity of *S. aureus* towards the spent media of the indicated *P. aeruginosa* strains was examined by quantifying their lag phase extensions (LPE). LPE were calculated as described before (49, 51) by determining the differences in time for spent-media-treated samples to reach *A*_600 nm_ > 0.45 compared to the untreated controls (i.e. *S. aureus* incubated with the sterile-filtered spent media of a 24 hr *S. aureus* culture). As additional controls, we also determined the sensitivities of the *P. aeruginosa* strains tested to *S. aureus* spent media. Here, overnight *P. aeruginosa* cultures were diluted into TSB to an OD_600_ of 0.02, incubated in the presence of the indicated ratios of sterile-filtered *S. aureus* spent media at 37 ^0^C and shaking, and OD_600_ measurements taken every 10 minutes for 16 hours in a Tecan Infinite 200 plate reader. LPE were calculated as described before.

### Competition assay in culture

Overnight cultures of *S. aureus* and the various *P. aeruginosa* strains were normalized an OD of 1 in fresh LB and cultivated independently and in combination as indicated. In competition, *S. aureus* and *P. aeruginosa* were cultivated in a 100:1 ratio. Samples were taken at the indicated time points, serial diluted, and plated onto cetrimide agar (Millipore Sigma) and DIFCO^TM^ mannitol salt agar (Ward’s Science) using the drip plate method (94) to count colony forming units (CFU) of *P. aeruginosa* and *S. aureus,* respectively.

### Quantification of polyP from the spent media of different *P. aeruginosa* strains

PolyP was quantified from the 24-hour old spent media of the different *P. aeruginosa* strains by using a modified version of the 4′,6-Diamidino-2-phenylindole dihydrochloride (DAPI)-PolyP assay (58). Here, 25 µM of DAPI (Sigma-Aldrich) was added to 500 µL of the spent media, incubated for 30 minutes in the dark and DAPI-polyP fluorescence was measured (exc. λ: 415 nm; em λ: 550 nm). The relative fluorescence units (RFUs) were calculated using sterile LB containing 25 µM DAPI as a control. PolyP concentrations were then extrapolated using a sodium polyP (Fisher Scientific) standard curve that was prepared in LB.

### Analysis of long-time survival in stationary phase *P. aeruginosa* strains

The indicated *P. aeruginosa* strains were grown in LB at 37 ^0^C and 300 rpm. Samples were taken at the indicated time points, serially diluted, and plated onto LB agar for CFU counts.

### Quantification of DNA from the spent media of different *P. aeruginosa* strains

DNA was quantified from the 24-hour old spent media of the different *P. aeruginosa* strains by DAPI (95). Here, 25 µM of DAPI (Sigma-Aldrich) was added to 500 µL of the spent media, incubated for 30 minutes in the dark, DAPI-DNA fluorescence measured (exc. λ: 350 nm; em λ: 470 nm). The relative fluorescence units (RFUs) were calculated using sterile LB containing 25 µM DAPI as a control.

### Analysis of membrane damage

The experiment was performed as previously described (96). Briefly, exponentially growing *S. aureus* cells were treated for one hour with the indicated concentrations of *(i)* sodium polyP (Fisher Scientific); *(ii)* purified pyocyanin (Cayman Chemicals); or *(iii)* the sterile-filtered spent media of the *P. aeruginosa* strains. 500 µL of each culture was pelleted, washed in PBS, and resuspended in 500 µL PBS containing 0.5 µM propidium iodide (PI). The samples were incubated in dark for 15 minutes and PI fluorescence (exc. λ: 535 nm; em λ: 615 nm) determined. The RFU values were normalized to the OD_600_ of the *S. aureus* culture.

### Determining the impact of polyP and pyocyanin on *S. aureus* growth and survival

Overnight TSB cultures of *S. aureus* were diluted into TSB to an OD_600_ of 0.03 and incubated in the presence and absence of the indicated concentrations of sodium polyP (Fisher Scientific) or pyocyanin (Cayman Chemicals) in a 96-well plate at 37 ^0^C under shaking conditions. PolyP was dissolved in ddH_2_O, purified pyocyanin (Cayman Chemicals) was stored in DMSO and working solutions were diluted into ddH_2_O. OD_600_ measurements taken every 10 minutes for 16 hours in a Tecan Infinite 200 plate reader and LPE as described before. For the survival assay, samples were taken after 24 hrs of incubation, serial diluted, and plated for CFU counts as described before.

### Quantification of Elastase activity

Elastase activity was quantified from the spent media of 24 hr old *P. aeruginosa* cultures using the Elastin Congo Red (ECR) assay previously described (97). Briefly, 50 µL spent media was incubated for 4 hrs in 1 mL of 20 mg ECR-containing Tris-maleate buffer at 37 ^0^C and 300 rpm, following which 100 µL of 0.12 M EDTA was added and samples centrifuged for 5 min at 11,000 x g. OD_495_ of the supernatant was measured and elastase activity calculated.

### Quantification of LasA protease activity

LasA protease activity was quantified from freshly prepared spent media of 24-hour *P. aeruginosa* cultures as previously described (10). Briefly, overnight *S. aureus* cells were heat-killed for 20 min at 95 ^0^C, cells spun down, and the cell pellet resuspended in 20 mM Tris-HCl (pH 8) to an OD_600_ of 1. The *P. aeruginosa* spent media were diluted 4-fold into LB broth and added to the heat-killed *S. aureus* cells. The samples were incubated at room temperature under static conditions and OD_600_ was measured at 0 (OD_initial_) and 2 (OD_final_) hours using a Tecan Infinite 200 plate reader. Lysis activity (i.e., LasA protease activity) was determined using the following formula:

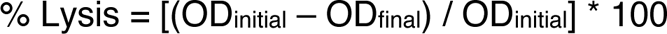

### Pyochelin quantification

Pyochelin was quantified from spent media of 24-hour *P. aeruginosa* cultures as previously described (98). Briefly, pyochelin was extracted twice from 1 mL spent media using 500 µL ethyl acetate. The OD_320_ of the extract was measured using a spectrophotometer and the pyochelin concentration was determined using ε_pch_ = 4,300 M^-1^ cm^-1^.

### Pyoverdine quantification

Pyoverdine was quantified from spent media of 24-hour *P. aeruginosa* cultures as previously described (99). Briefly, pyoverdine was quantified from the spent media fluorometrically (exc. λ: 400 nm; em λ: 477 nm) using the Tecan Infinite 200 plate reader. The fluorescence readings were normalized to the OD_600_ of the respective culture.

### Rhamnolipid quantification

Rhamnolipid was quantified from spent media of 24-hour *P. aeruginosa* cultures as previously described (100). Briefly, 40 µL of 1N HCl was used to acidify 1 mL of the spent media, following which the rhamnolipids were extracted using 4 mL chloroform. 100 µL of 1 mg/mL methylene blue was added to 3 mL of chloroform extract and samples vortexted and centrifuged after addition of 4 mL ddH_2_O. OD_638_ of the chloroform layer was measured to quantify rhamnolipids.

### Pyocyanin quantification

Pyocyanin was quantified from spent media of 24-hour *P. aeruginosa* cultures as previously described (101). Briefly, 750 µL of chloroform was used to extract pyocyanin from 1.3 mL of spent media. 600 µL of the resulting blue layer was acidified with 300 µL of 0.2 N HCl, resulting in a color change to pink. OD_520_ of the pink layer was determined using the Tecan Infinite 200 plate reader and multiplied by 17.062 to obtain the amount of pyocyanin in µg/mL.

### Determining the antimicrobial effects of pyocyanin

Overnight cultures of *S. aureus* were diluted into fresh TSB to an OD_600_ of 0.05 and incubated in the presence and absence of the indicated concentrations of pyocyanin at 37 ^0^C under shaking conditions. Samples were taken after 23 hrs of incubation, serial diluted, and plated for CFU counts.

### Determining *S. aureus* sensitivity to hydrogen peroxide using a growth assay

Overnight cultures of *S. aureus* strain *USA300JE2ΔkatAΔahpC* were diluted into TSB to an OD_600_ of 0.03 and cultivated in the presence and absence of hydrogen peroxide (Sigma-Aldrich) at 37 ^0^C and under shaking conditions in a Tecan Infinite 200 plate reader. OD_600_ measurements were taken every 10 minutes for 16 hours.

### Analysis of *katB* transcript level using qRT-PCR

Exponentially growing *S. aureus* USA300LAC*ΔkatAΔahpC* cells were treated with 40 µg/mL pyocyanin or the indicated spent media of 24 hr *P. aeruginosa* cultures in a 3:1 (v/v) ratio for 10 minutes, following which ice-cold methanol was added to stop transcription. Cell pellets were lysed with 100 mg/mL lysostaphin, RNA extracted using the RNA Extraction Kit (Macherey & Nagel), residual genomic DNA removed using the TURBO DNA-Free Kit (Thermo Scientific), and mRNA reverse-transcribed into cDNA using the PrimeScript cDNA Synthesis Kit (TaKaRa). qRT-PCR was set up according to the manufacturer’s instructions (Alkali Scientific) using the following primers: *katB-fw*, CGAGGATTTGCGTTAAAGTTC; *katB-rev*, ACCGCGCGATTTAAACTAAC; *rrlB-fw,* TTTAGCCCCGGTACATTTTCG; *rrlB-rev,* TTTAGCCCCGGTACATTTTCG. The *katB* transcript level were normalized to the transcript level of the 23S rRNA-encoding *rrlB* gene. The relative fold-changes in gene expression were calculated using the ΔΔCt method (102).

## ACKNOWLEDGEMENTS

This work was supported by the NIAID grants R15AI164585 and 1R03AI174033-01A1 and the Illinois State University Faculty Research Award (to J.-U. D.). R.S. was supported by a Weigel fellowship by the Phi-Sigma Biological Sciences Honors Society. We thank Dr. Thomas Kehl-Fie (University of Illinois Urbana-Champaign) for providing the USA300 JE2 wildtype and *ΔahpCΔkatA* strains. Members of the Dahl lab are acknowledged for feedback and proof-reading of the manuscript. Figure 6 was created using Biorender.

## Notes

### Competing Interest Statement

The authors have declared no competing interest.

