## Supplemental Fig S1-5 for "Pseudomonas aeruginosa kills Staphylococcus aureus in a polyphosphate-dependent manner"

### SUPPLEMENTARY INFORMATION

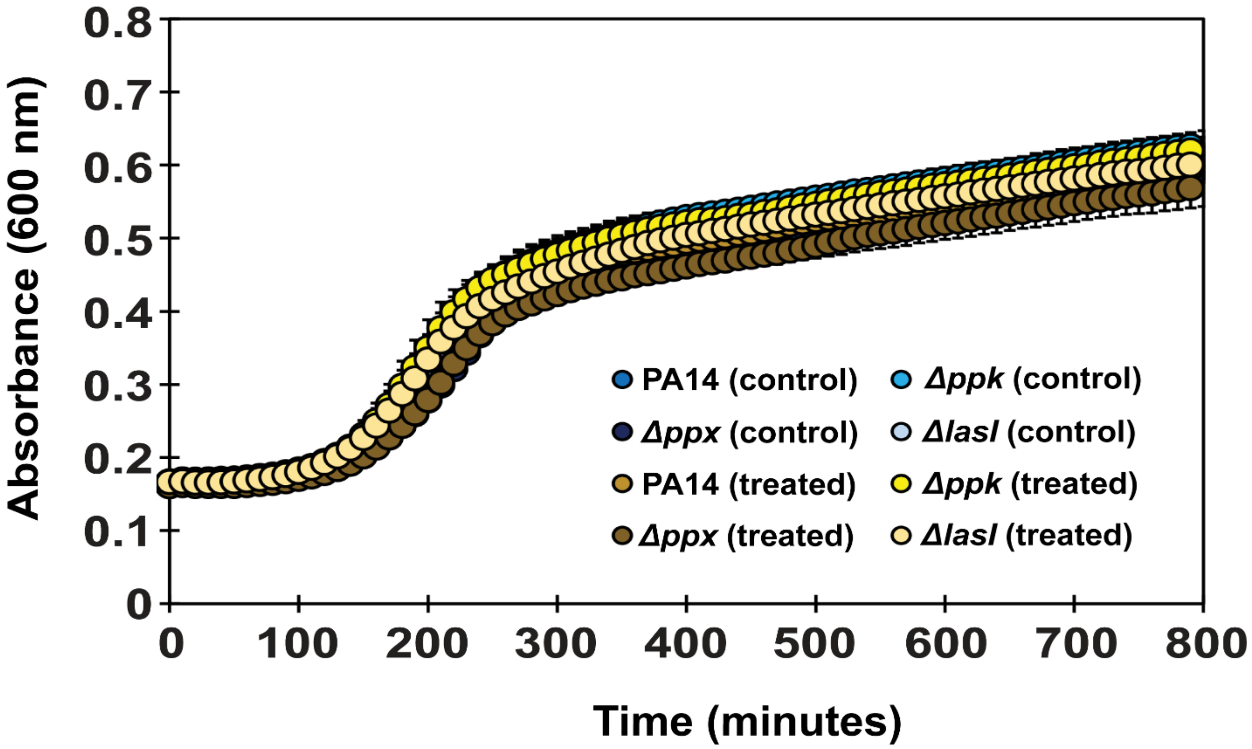


**Supplementary FIG S1. *P. aeruginosa* is not inhibited by *S. aureus* spent media.** The indicated *P. aeruginosa* strains were diluted into TSB to an $OD_{600}$ of 0.03 and exposed to the spent media of a 24 hrs, stationary phase *S. aureus* culture in a 12.5:1 ratio. $A_{600nm}$ was measured every 10 minutes for 16 hours using the Tecan Infinite 200 plate reader *(n = 3 [with 4 technical replicates], ± S.D.)*.

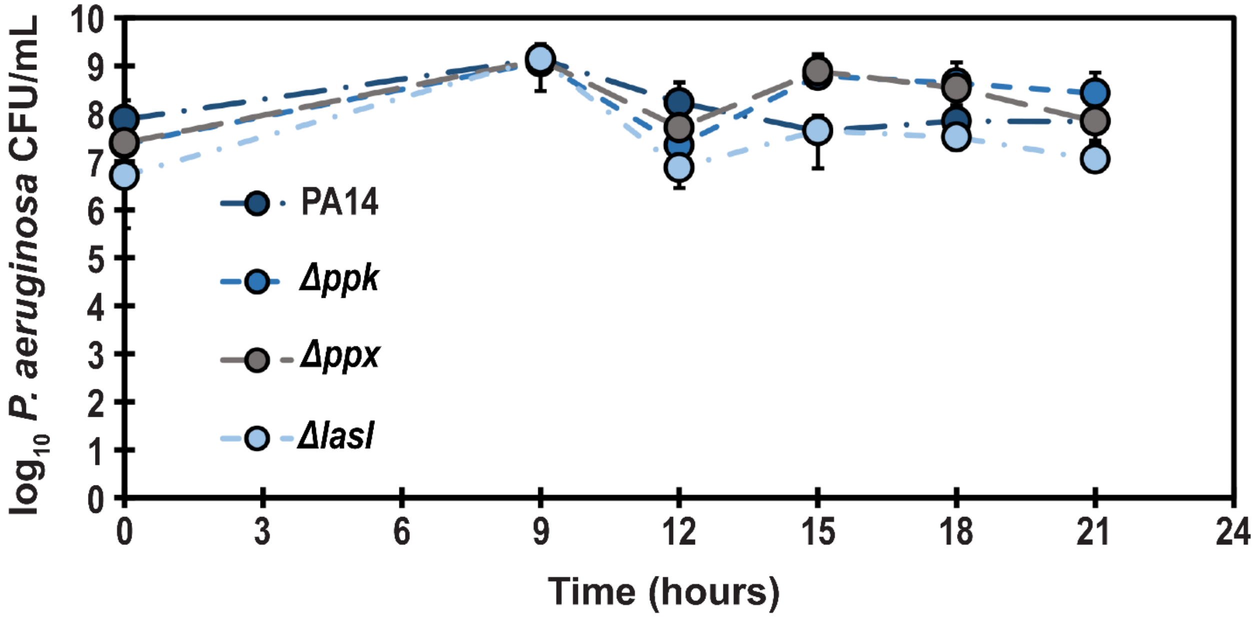


**Supplementary FIG S2. *P. aeruginosa* survival is not affected by *S. aureus* in a co-cultivation assay.** Overnight cultures of *S. aureus* strain USA300LAC and the indicated *P. aeruginosa* strains were each diluted into TSB media to an $OD_{600}$ of 1, mixed in a ratio of 100:1, and cultivated at 37 °C for 21 hrs. Samples were taken at indicated time points and serially diluted for *P. aeruginosa* CFU counts on cetrimide agar. Dark blue circles, PA14; blue circles, Δ*ppk*; bright blue circles, Δ*lasI;* grey circles, Δ*ppx; (n = 3, ± S.D.).*

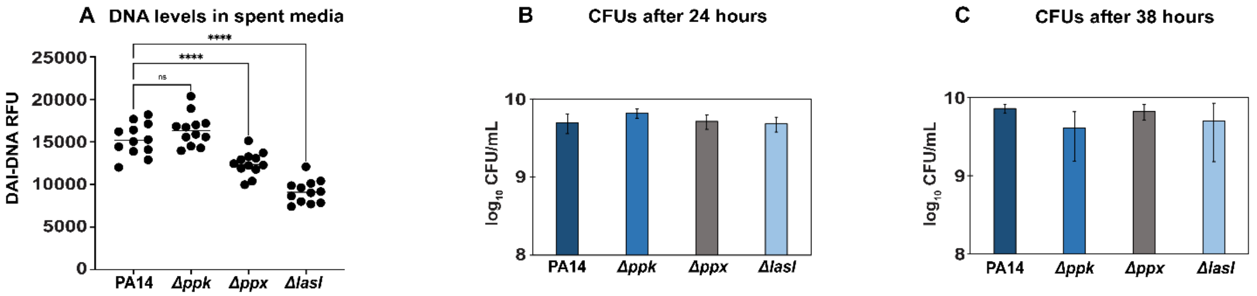


**Supplementary FIG S3. *P. aeruginosa* long-term survival is not affected by polyP production. (A)** Extracellular DNA levels were determined in the spent media of 24 hrs, stationary phase *P. aeruginosa* cultures using 25 μM DAPI. DAPI-DNA fluorescence was measured at excitation/emission wavelengths of 350 and 470 nm, respectively; *(n = 8, ± S.D.).* **(B & C)** Samples were taken from the indicated *P. aeruginosa* cultures after **(B)** 24 hrs and **(C)** 38 hrs of cultivation in LB at 37 °C and 300 rpm, serially diluted and plated onto LB agar for CFU counts; *(n = 3-4, ± S.D.)*.

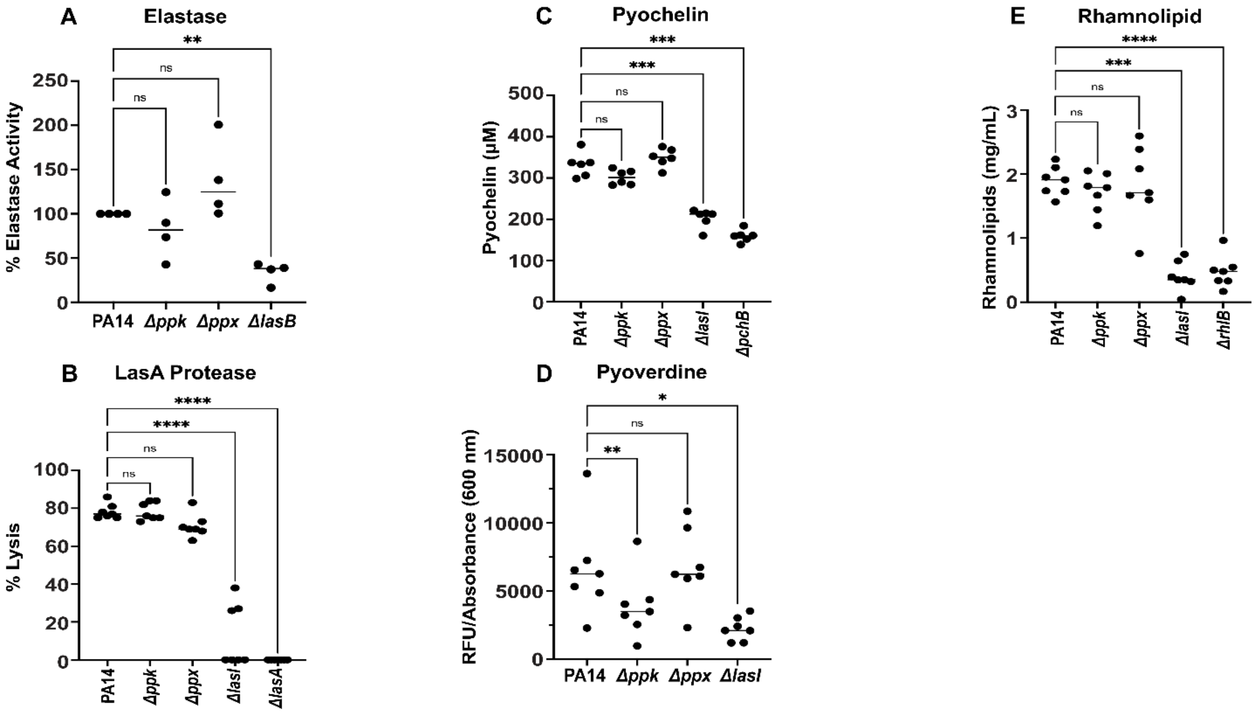


**Supplementary FIG S4. The impact of polyP metabolism on *P. aeruginosa* virulence factors.** The following virulence factors were quantified from sterile-filtered spent media of the indicated 24 hrs stationary phase *P. aeruginosa* cultures as described in *Material and Methods*: **(A)** Elastase *(n = 5, ± S.D.)*; **(B)** LasA Protease *(n = 7, ± S.D.)*; **(C)** Pyochelin *(n = 6, ± S.D.)*; **(D)** Pyoverdine *(n = 6, ± S.D.)*; and **(E)** Rhamnolipids *(n = 7, ± S.D.)*. *Statistical tests: One-Way ANOVA, ns: p > 0.05, * p ≤ 0.05, ** p ≤ 0.01, *** p ≤ 0.001, **** p ≤ 0.0001).*

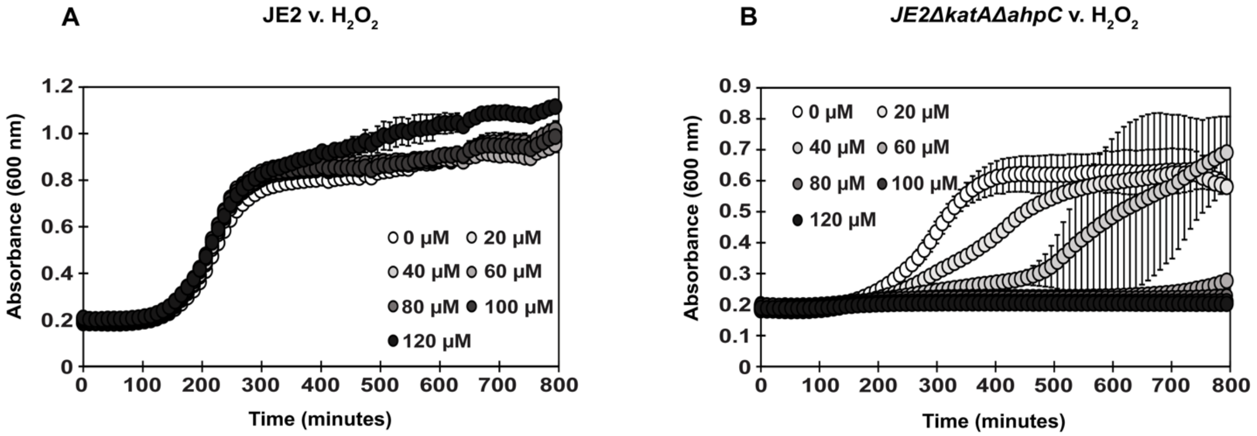


**Supplementary FIG S5. Lag phase extension assays of different *S. aureus* strains.** *S. aureus* strains reader **(A)** USA300LAC and **(B)** USA300LAC*ΔkatAΔahpC* were diluted into TSB to an $OD_{600}$ of 0.03 and cultivated with and without the indicated concentrations of $H_2O_2$. $A_{600nm}$ was measured every 10 minutes for 16 hours using the Tecan Infinite 200 plate reader, *(n = 3 [with 2 technical replicates], ± S.D.)*
